# 4β-hydroxycholesterol is a pro-lipogenic factor that promotes SREBP1c expression and activity through Liver X-receptor

**DOI:** 10.1101/2020.08.20.256487

**Authors:** Ofer Moldavski, Peter-James H. Zushin, Charles A. Berdan, Robert J. Van Eijkeren, Xuntian Jiang, Mingxing Qian, Daniel S. Ory, Douglas F. Covey, Daniel K. Nomura, Andreas Stahl, Ethan J. Weiss, Roberto Zoncu

## Abstract

Oxysterols are oxidized derivatives of cholesterol that play signaling roles in lipid biosynthesis and homeostasis. Here we show that 4β-hydroxycholesterol (4β-HC), a liver and serum abundant oxysterol of poorly defined function, is a potent and selective inducer of the master lipogenic transcription factor, Sterol Regulatory Element Binding Protein 1c (SREBP1c), but not the related steroidogenic transcription factor SREBP2. Mechanistically, 4β-HC acts as a putative agonist for Liver X receptor (LXR), a sterol sensor and transcriptional regulator previously linked to SREBP1c activation. Unique among the oxysterol agonists of LXR, 4β-HC induced expression of the lipogenic program downstream of SREBP1c, and triggered *de novo* lipogenesis both in primary hepatocytes and in mouse liver. 4β-HC-acted in parallel to insulin-PI3K-dependent signaling to stimulate triglyceride synthesis and lipid droplet accumulation. Thus, 4β-HC is an endogenous regulator of de novo lipogenesis through the LXR-SREBP1c axis.

## INTRODUCTION

All cells must achieve and maintain a balanced composition of their internal membranes in order to grow, proliferate or adapt to sudden changes in external conditions and nutrient availability (Nohturfft and Zhang, 2009). Dedicated biosynthetic pathways mediate the synthesis of fatty acids, sterols, phospholipids and sphingolipids, but how these pathways communicate with each other to coordinate their respective activities and respond to changing metabolic needs is poorly understood (Thelen and Zoncu, 2017; van Meer et al., 2008).

Liver-X-Receptor (LXR) α and β are transcription factors belonging to the nuclear receptor superfamily that play key roles in maintaining lipid homeostasis in multiple cells and organs (Janowski et al., 1996; Kalaany and Mangelsdorf, 2006; Repa et al., 2000; Yoshikawa et al., 2001). LXR α and β dimerize with the retinoic X-receptor (RXR) and activate target genes that mediate cholesterol efflux from cells, including ABC-family transporters, as well as genes that mediate conversion of cholesterol into bile acids in the liver to facilitate cholesterol elimination from the body, such as cytochrome p450 7a-hydroxylase (CYP7A1) (Costet et al., 2000; Peet et al., 1998; Svensson et al., 2003). Accordingly, mice lacking LXRα exhibit impaired bile acid metabolism and defective cholesterol elimination (Peet et al., 1998), along with enhanced inflammation and formation of atherosclerotic plaques (Hong et al., 2012). Conversely, synthetic LXRα agonists have shown promise in reducing atherosclerosis and preventing cardiovascular disease in animal models (Calkin and Tontonoz, 2010; Joseph et al., 2002; Terasaka et al., 2003).

Another key mediator of lipid homeostasis is the helix-loop-helix-leucine zipper transcription factor, Sterol Regulatory Element Binding Protein (SREBP) 1c. SREBP1c is a master regulator of biosynthesis of fatty acids and triglycerides (collectively referred to as *de-novo* lipogenesis, DNL) that is subject to tight transcriptional and post-translational regulation. Along with its paralogue, the master steroidogenic transcription factor SREBP2, SREBP1c resides at the endoplasmic reticulum (ER) membrane, to which it is anchored via a single transmembrane helix. When cholesterol concentration in the ER membrane is low, SREBP1c and SREBP2 are transported to the Golgi apparatus via interaction with SREBP cleavage-activating protein (SCAP), a cholesterol-sensing chaperone that favors their loading into COPII vesicles. At the Golgi membrane, resident proteases cleave the DNA-binding portion of SREBP1c and SREBP2 from the transmembrane portion, enabling their translocation to the nucleus and activation of downstream programs for *de-novo* lipogenesis and steroidogenesis, respectively.

In addition to their homeostatic regulation by cholesterol levels, the SREBPs lie downstream of metabolic hormone signaling. For example in the liver, both the expression and proteolytic activation of SREBP1c are stimulated by the insulin-phosphatidylinositol 3-kinase (PI3K)-mechanistic Target of Rapamycin (mTOR) pathway, as part of a mechanism that converts excess of glucose into lipids, which are required for energy storage (Azzout-Marniche et al., 2000; Horton et al., 2002; Ricoult and Manning, 2013). However, the range of regulatory inputs to SREBP1c and their respective interplay remain to be fully elucidated.

LXRα and LXRβ were shown to directly bind to the promoter of the SREBP1c gene and trigger activation of its downstream lipogenic genes (Repa et al., 2000). Accordingly, synthetic LXR ligands strongly promote de novo lipogenesis and increased plasma triglyceride levels (Grefhorst et al., 2002; Joseph et al., 2002; Schultz et al., 2000), providing evidence for cross-talk between LXR- and SREBP1c-dependent programs.

While the physiological significance of LXR-dependent regulation of DNL through SREBP1c remains unclear, this cross-talk has important clinical implications. In particular, LXR-dependent upregulation of SREBP1c potentially limits the usefulness of LXR agonists to improve cholesterol metabolism, as the resulting induction of lipogenic programs could lead to undesirable effects, such as non alcoholic fatty liver disease (NAFLD), a condition that has risen to epidemic proportions in recent years (Cai et al., 2018). Thus, understanding how LXR-dependent activation of SREBP1c occurs, and its functional interaction with other pathways controlling lipid homeostasis such as PI3K-mTOR signaling are key open questions.

Oxysterols are a family of metabolites that originate from an oxygenation reaction of cholesterol. Some Oxysterols are signaling molecules involved in a wide range of physiological processes controlling cholesterol, glucose and lipid metabolism (Mutemberezi et al., 2016). Levels of oxysterols are known to change in pathological situations like obesity, atherosclerosis and Alzheimer disease (Guillemot-Legris et al., 2016b; Poli et al., 2013). A subset of oxysterols function as endogenous LXR ligands and were shown to activate LXRα-dependent gene expression *in vitro*, including those bearing hydroxyl groups in position 4, 7, 20, 22, 24, 25 and 27 on the cholesterol backbone (Janowski et al., 1999; Janowski et al., 1996; Nury et al., 2014). Interestingly, although these oxysterols are considered *bona fide* LXR activators, none is known to activate SREBP1c and its downstream lipogenic programs, whereas several oxysterols have been shown to promote LXR-dependent cholesterol efflux. In contrast, synthetic LXR ligands including T0901317 and GW3965 can induce both cholesterol efflux and SREBP1c-dependent DNL (Grefhorst et al., 2002; Schultz et al., 2000). This leads to the question of whether *de novo* lipogenesis is a physiologically relevant LXR-dependent response, and if so, the identity of the endogenous ligand that triggers LXR-dependent SREBP1c expression.

Here we identify 4β-hydroxycholesterol (4β-HC) as an LXR activator that selectively triggers SREBP1c activation and *de novo* fatty acid and triglyceride synthesis. 4β-HC promoted the expression and proteolytic processing of SREBP1c but not of the related steroidogenic factor SREBP2, thus triggering de novo synthesis of fatty acids but not cholesterol. In primary mouse hepatocytes, 4β-HC additively enhance insulin action in promoting SREBP1c expression and activation, leading to increased triglyceride synthesis and storage. Thus, 4β-HC may be a novel lipogenic factor that can shift lipid homeostasis towards triglycerides accumulation via regulation on SREBP1c.

## RESULTS

### 4β-HC is a unique oxysterol that drives SREBP1c gene expression

To identify oxysterol ligands that could promote SREBP1c expression, we treated liver carcinoma-derived Huh7 cells with a panel of oxysterols selected among the most abundant in the bloodstream, including 4β-, 7β-, 19-, 20-, 24(S)-, 25- and 27-hydroxycholesterol (HC). By quantitative PCR (qPCR), several oxysterols previously identified as LXR activators, including 4β-HC, 7β-HC, 24(S)-HC and 25-HC, induced the expression of a canonical LXR target gene, ATP binding cassette subfamily A member 1 (ABCA1), with variable potency (Fig. 1A). In contrast, 4β-HC was the only oxysterol to induce significant upregulation of the SREBP1c transcript (Fig. 1B). A dose-response comparison between 4β-HC and 24(S)-HC showed that 24(S)-HC is a more potent activator than 4β-HC toward ABCA1 (Fig. 1C) and another canonical LXR gene target, ABCG1 (Fig. 1D). Conversely, 4β-HC activated SREBP1c, more potently than 24(S)-HC (Fig. 1E). 4β-HC-mediated induction of the SREBP1c gene was enantioselective, as the non-natural enantiomer of 4β-HC (ent-4HC) was unable to induce SREBP1c mRNA expression even at the highest concentration used (20μM) (Fig. 1F). This data suggest that SREBP1c induction depends on unique structural features of 4β-HC.

**Figure 1:**
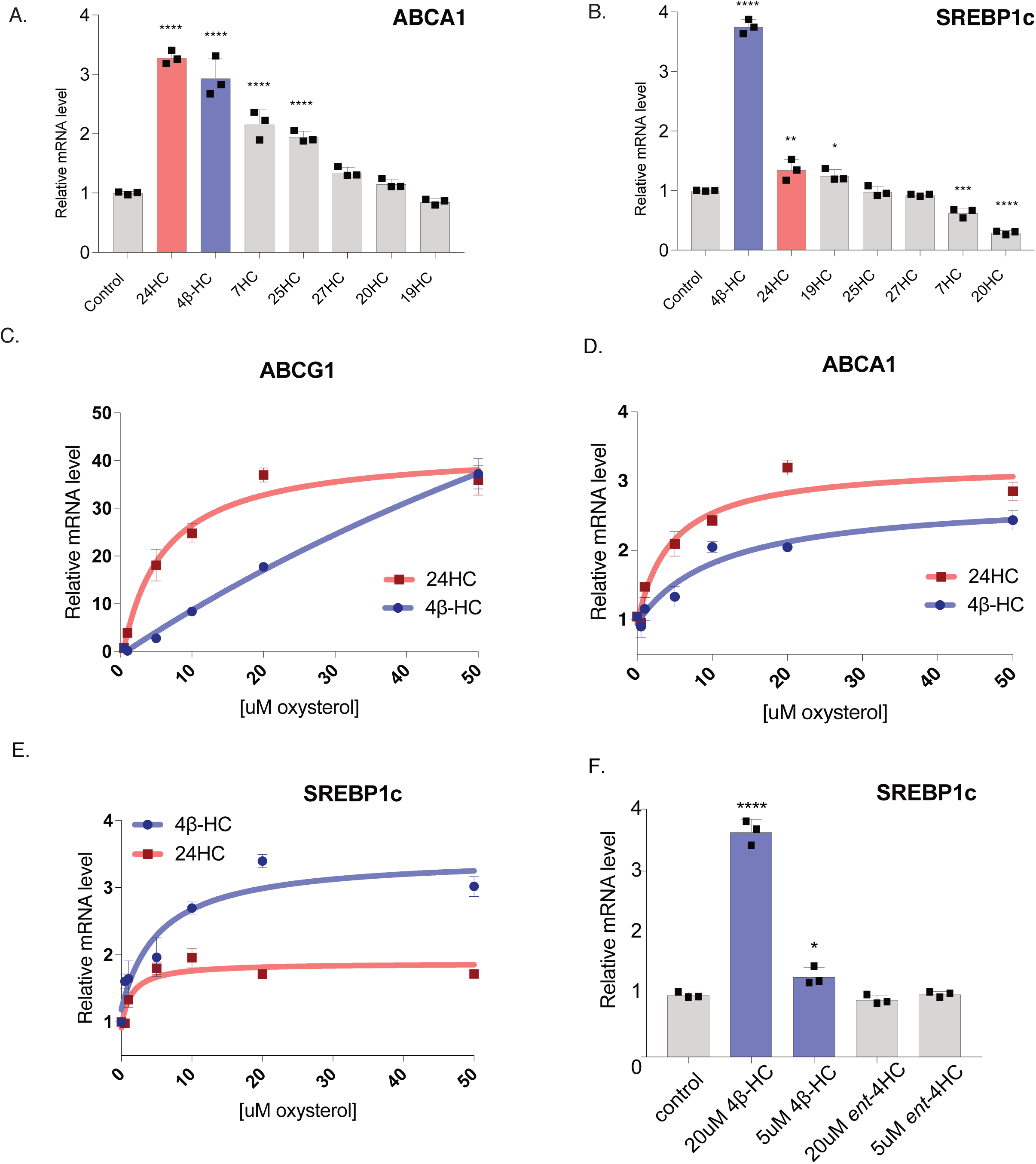
The oxysterol 4β-HC selectively upregulates SREBP1c expression. **A**. Oxysterol screen for LXR target gene expression. Huh7 cells were treated with indicated oxysterols (20μM) in 24 hours time course. ABCA1 mRNA levels or (**B**) SREBP1c mRNA level were measured by RT-PCR. **C**. Dose response curves of 4β-HC and 24HC in Huh7 cells treated for 24 hours. mRNA levels of ABCA1, (**D)** ABCG1 and (**E)** SREBP1c were measured by RT-PCR. Line plotted by non-linear fit. **E**. SREBP1c induction by 4β-HC is stereo-specific. Huh7 cells were treated with 4β-HC or an enantiomer of 4HC (ent-4HC) for 24 hours in the indicated concentration. Bars are Mean+SD. Statistical significance calculated by one-way ANOVA. *p<0.05, **p<0.01, ***p<0.001, ****p<0.0001, NS: Not significant.

### 4β-HC induces expression and activation of SREBP1 but not SREBP2

Oxysterols such as 25- and 27-hydroxycholesterol suppress SREBP1 and SREBP2 activation by blocking their trafficking to the Golgi, where proteolytic processing of the SREBPs to the mature nuclear form occurs (Adams et al., 2004; Radhakrishnan et al., 2007).

In contrast to these oxysterols, 4β-HC significantly increased SREBP1c mRNA levels (Fig 2A) as well as protein levels in a cycloheximide-sensitive manner (Fig 2B). However, 4β-HC did not increase either mRNA or protein levels of SREBP2 (Fig 2A-2B). In keeping with the increased total levels of SREBP1c, 4β-HC increases both cytosolic and nuclear forms of SREBP1 in a dose-dependent manner, whereas levels of cytoplasmic or nuclear SREBP2 protein levels did not change (Fig 2C). Consistent with previous reports (Adams et al., 2004; Radhakrishnan et al., 2007) and in contrast to 4β-HC, 25-HC reduced the nuclear forms of both SREBP1 and SREBP2, thereby causing the accumulation of the unprocessed cytoplasmic form of both proteins but without transcriptional upregulation (Fig. 1B).

**Figure 2:**
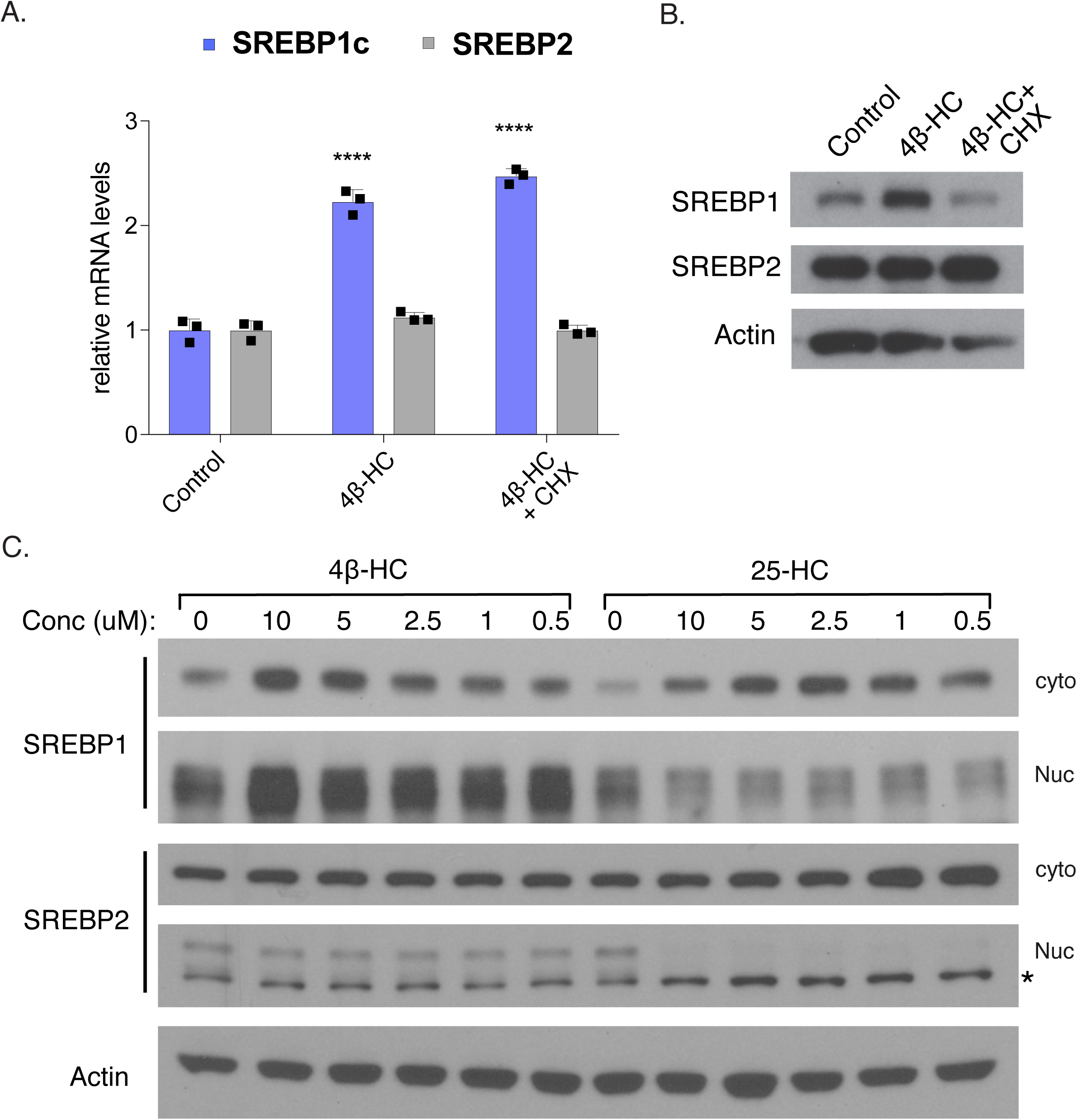
4β-HC induces expression and activation of SREBP1 but not SREBP2. **A**. 4β-HC increases SREBP1 protein expression. Huh7 cells were treated with 20μM 4β-HC and a translation inhibitor, cycloheximide (CHX) for 4 hours followed by measurement of SREBP1 and SREBP2 mRNA and **(B**.) protein level. **C**. 4β-HC increases SREBP1 cytosolic and nuclear levels while not affecting SREBP2. Huh7 cells were treated with 4β-HC or 25-HC for 24hours followed cytosolic-nuclear fractionation to measure protein level of SREBP1 and SREBP2 cytoplasmic and nuclear levels. Cyto: Cytosolic. Nuc: Nuclear. Asterisk denotes unspecific band in SREBP2 nuclear blot. Bars are Mean+SD. Statistical significance calculated by one-way ANOVA. ****p<0.0001.

This data suggest that, unlike other oxysterols that function as inhibitors of both SREBP1c and SREBP2, 4β-HC is a specific inducer of SREBP1c expression and activation.

### 4β-HC induce lipogenic programs through the LXRs

Along with other oxysterols, 4β-HC was previously shown to activate LXRα-dependent transcription in luciferase assays *in vitro*, supporting its role as a putative LXR ligand (Janowski et al., 1996; Nury et al., 2013). In turn, LXR transcriptionally activates SREBP1c by directly binding to its promoter region (Repa et al., 2000). Combining these observations, we thus hypothesized that 4β-HC may transcriptionally activate SREBP1c and its downstream lipogenic programs via LXR. Consistent with this possibility, co-treating cells with 4β-HC together with an LXR antagonist (GSK-2033) abolished 4β-HC-dependent induction of SREBP1c gene expression (Fig. 3A).

**Figure 3:**
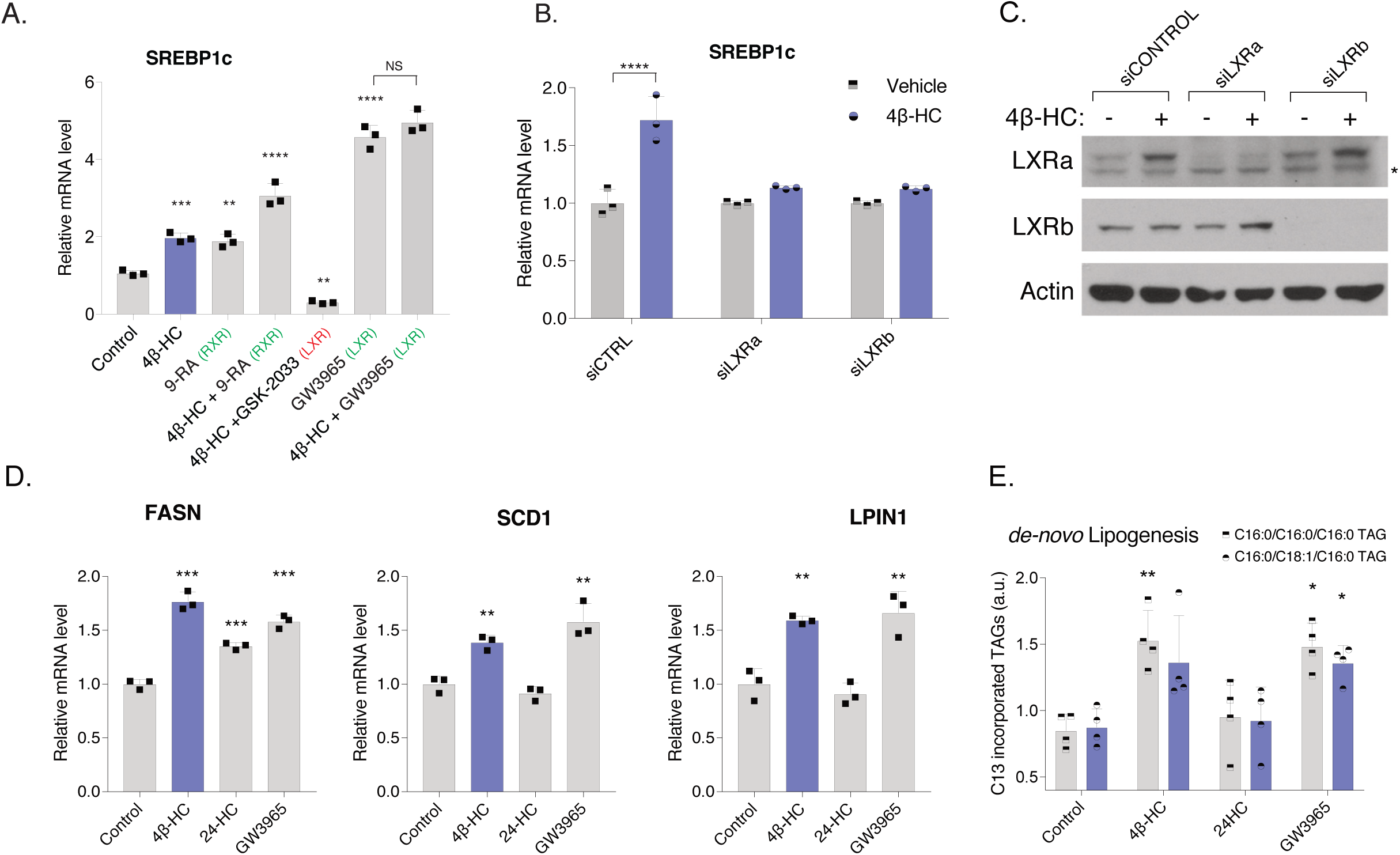
4β-HC induce lipogenic programs through the LXRs. **A**. 4β-HC interacts with LXR and RXR agonist and antagonists like an LXR ligand. Huh7 cells were treat with 20μM 4β-HC, RXR agonist, 9-cis-retinoic acid (9-RA), LXR antagonist (GSK-2033) and LXR agonist (GW3965). For convenience, Agonists are marked in green and antagonist are marked in red. **B**. LXRα and LXRβ are required for SREBP1c induction by 4β-HC in Huh7 cells. Knockdown of LXRα or LXRβ by siRNA for 72 hours followed by treatment with 5μM 4β-HC for 24 hours followed by RT-PCR of SREBP1c. **C**. Knockdown efficiency was evaluated by measurement of LXRα and LXRβ protein levels **D**. 4β-HC induction of lipogenic genes. Huh7 cell were treat for 24 hours with 4β-HC, 24-HC or LXR agonist (GW3965) followed by mRNA measurement of Fatty acid synthase (FASN), Stearoyl CoA Desaturase 1 (SCD1) and Lipin1 (LPIN1). **E**. 4β-HC increases *de-novo* Lipogenesis. Huh7 treated for 24 hours with 5μM 4β-HC, 24-HC or LXR agonist (GW3965) with media containing C13 Glucose followed by Lipids extraction. C13 incorporation into TAGs was measured via Liquid chromatography Mass spec. Asterisk denotes unspecific band in LXRα blot. Bars are Mean+SD. Statistical significance calculated by one-way ANOVA. *p<0.05, **p<0.01, ****p<0.0001.

The effect of 4β-HC on SREBP1c induction was additive with an RXR ligand, 9-cis Retinoic acid (9-RA). Moreover, co-incubation of 4β-HC with the LXR agonist, GW3965, used at concentrations that activate LXR maximally, caused no additional increase in SREBP1c expression over GW3965 alone (Fig. 3A). siRNA-mediated knock down of either LXR*α* or LXRβ (both of which are expressed in Huh7 cells) largely abolished 4β-HC-dependent SREBP1c mRNA expression (Fig. 3B). Interestingly, we noticed that 4β-HC treatment increased LXR*α* protein levels, a stabilizing effect observed for other established LXR ligands (Ignatova et al., 2013) (Fig. 3C). Together, and combined with previous reports these data support the hypothesis that 4β-HC induces SREBP1c gene expression by acting as an LXR agonist.

We next compared the ability of 4β-HC to induce SREBP1c-dependent lipogenic programs with that of the LXR agonist, GW3965. Fatty acid synthase (FASN), Stearoyl-CoA desaturase (SCD1) and Lipin1 (LPIN1) are validated SREBP1c downstream targets in Huh7 cells (Ishimoto et al., 2009; Shimomura et al., 1998). Treatment with either 4β-HC or GW3965 significantly increases the expression of these genes (Fig 3D). In contrast, 24-HC, another putative LXR ligand that failed to induce SREBP1c in our hands (Fig. 1B and 1E), had minimal or no effect on these SREBP1c target genes (Fig. 3D).

Previous work had shown that GW3965 induces FASN to a greater extent than the 1.6-fold we observed in Huh7 (Peng et al., 2011). Huh7, a hepatocellular carcinoma line, are known to hyperactivated DNL to supply membranal lipids required for rapid division and growth (Calvisi et al., 2011; Li et al., 2016). We speculate that the modest increase in FASN by GW3965 or 4β-HC is due to already elevated baseline expression that cannot be increased much further. To further substantiate the pro-lipogenic effect of 4β-HC, we directly measured DNL by C13 incorporation into triglycerides using liquid chromatography coupled to mass spectrometry (LC/MS). Similarly to lipogenic gene induction, both GW3965 and 4β-HC had a modest but statistically significant 1.5-fold increase in C13-labeled C16:C16:C16 TAG, or trending toward significance for C16:C18:C16 TAG, whereas 24-HC caused no significant change (Fig 3E)-Combined, these data suggest that the pro-lipogenic action of 4β-HC is comparable, in mechanism and potency, to known LXR agonists.

### 4β-HC induces lipid droplet formation and triglyceride accumulation

In keeping with the ability of 4β-HC to upregulate fatty acid biosynthetic genes via SREBP1c, treating Huh7 cells with 4β-HC (but not with its unnatural enantiomer, *ent*-4HC) for 72 hours resulted in marked accumulation of lipid droplets (LDs), as revealed by staining with the lipophilic dye BODIPY 493/503 (Fig. 4A and 4B). LD accumulation induced by 4β-HC was suppressed by simultaneous treatment with a FASN inhibitor, TVB-3166, or with the LXR inhibitor GSK-2033. Measurement of triglyceride content in cell extracts confirmed the ability of 4β-HC to induce triglyceride accumulation, albeit with lower potency than the LXR agonist GW3965, whereas cholesterol levels remained unchanged (Fig. 4C). Consistent with the BODIPY staining, both LXR and FASN inhibitors hindered 4β-HC-induced triglyceride accumulation (Fig. 4C). Moreover, as seen with SREBP1c induction, the enantiomer of 4β-HC (*ent*-4HC) failed to induce triglyceride accumulation (Fig. 4C). Thus, 4β-HC is sufficient to induce the formation of triglyceride-containing lipid droplets in an LXR- and FASN-dependent manner in cell culture.

**Figure 4:**
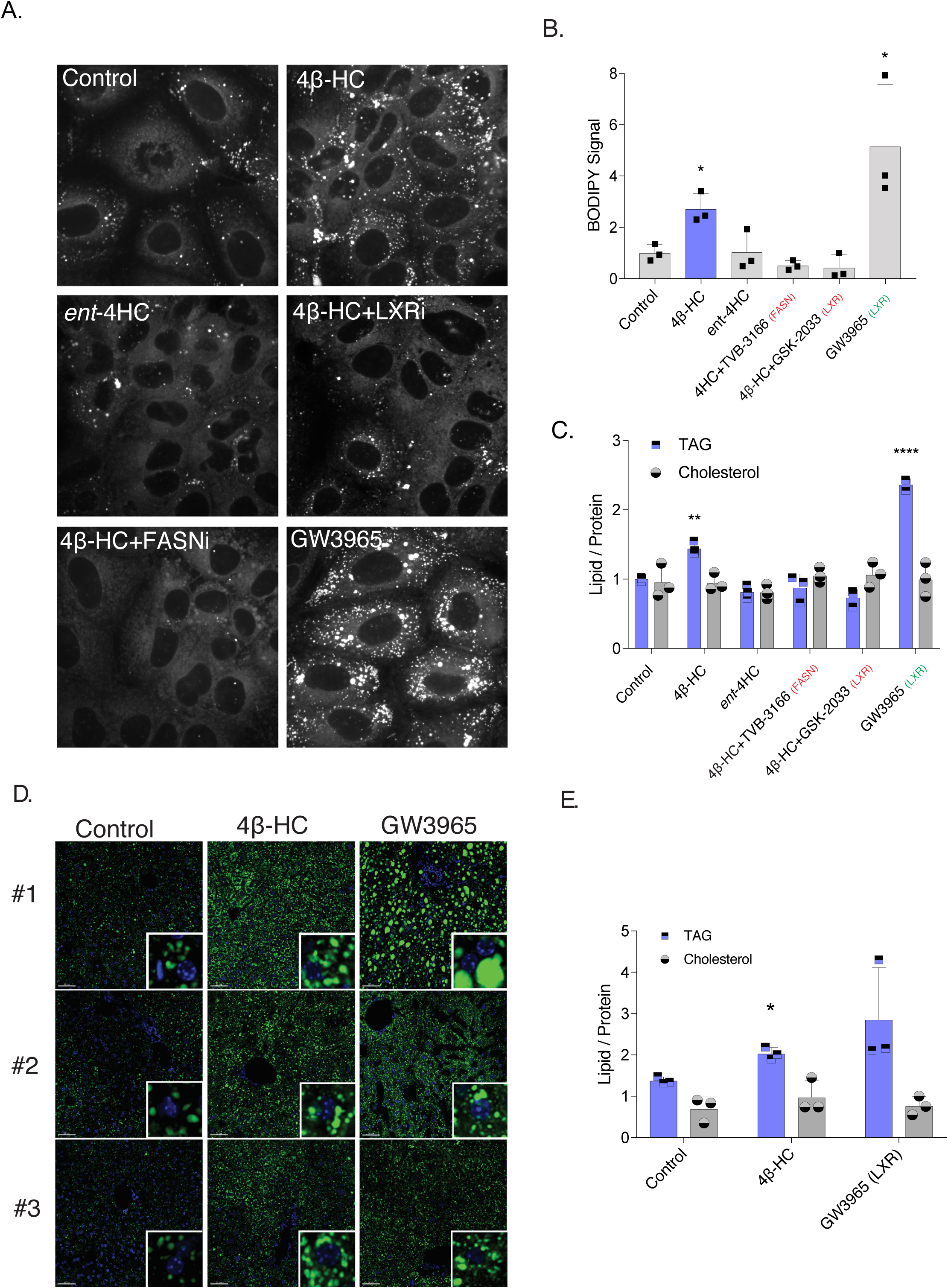
4β-HC induces lipid droplet formation and triglyceride accumulation. **A**. 4β-HC increases Lipid droplet size and number. Huh7 cells were treated with 5μM 4β-HC with indicated drugs for 72 hours followed staining with lipid droplet dye, BODIPY 493/503 and visualization by confocal microscopy and (**B**.) quantified using ImageJ. **C**. 4β-HC increases triglycerides (TAG) levels. Huh7 cells were treat as (C) followed by measurement of triglycerides, total cholesterol and protein levels using commercial kits. **D**. 4β-HC increase lipid droplet in mice liver. Mice were fed normal chew with either Vehicle, 50mg/kg/day 4β-HC or 10mg/kg/day GW3965 for 5 days. Liver samples from were fixed and stained with BODIPY 493/503 and DAPI to observe lipid droplet and nuclei ultrastructure. **E**. 4β-HC increases triglycerides (TAG) levels in mice liver treat above, followed by measurement of triglycerides, total cholesterol and protein levels using commercial kits. For convenience, Agonist are marked in green and antagonist are marked in red. Bars are Mean+SD. Statistical significance calculated by one-way ANOVA. ent-4HC: stereo enantiomer-4HC, FASNi: TVB-3166 LXRi: GSK-2033, *p<0.05, **p<0.01, ***p<0.001, ****p<0.0001.

Next, we tested the effect of 4β-HC on *in-vivo* lipogenesis by feeding mice a normal diet supplemented with either 4β-HC or GW3965. After 7 days, livers were harvested, lipid droplets were assessed by BODIPY staining, and liver lipid content (normalized to protein mass) was measured. Consistent with the results in Huh7 cells, 4β-HC significantly increased the size and number of lipid droplets in liver sections (Fig 4D) and liver triglyceride content (Fig 4E), albeit with lower potency than the synthetic LXR agonist, GW3965. Collectively, these data suggest that 4β-HC is a pro-lipogenic factor that can increase liver lipid content *in vivo*.

### 4β-HC acts in parallel to insulin-PI3K signaling to drive SREBP1c expression

Insulin is a key hormone that drives SREBP1c transcription, proteolytic processing and DNL in the postprandial state. Insulin regulates SREBP1c transcription via poorly understood mechanisms, which include AKT-dependent transcriptional downregulation of Insig-2a, the ER-retention factor that blocks translocation of SCAP-SREBP1c to the Golgi (Yabe et al., 2003; Yecies et al., 2011). LXR was shown to be required for insulin-dependent activation on SREBP1c (Chen et al., 2004), but whether and how insulin activates LXR is not understood.

To interrogate the relationship between 4β-HC and insulin signaling in driving SREBP1c transcription and processing, we used an insulin-responsive primary mouse hepatocytes (Foretz et al., 1999). In these cells, stimulation with either 4β-HC or insulin alone increased the mRNA levels of SREBP1c, while combined 4β-HC and insulin increased SREBP1c mRNA levels additively [as previously shown for LXR agonists (Chen et al., 2004)] (Fig. 5A). Interestingly, treatment with PI3K or mTORC1 inhibitors abolished SREBP1c induction by both Insulin and 4β-HC (Fig. 5A), raising the possibility that 4β-HC may act in a common pathway with insulin-PI3K-mTORC1 signaling.

**Figure 5:**
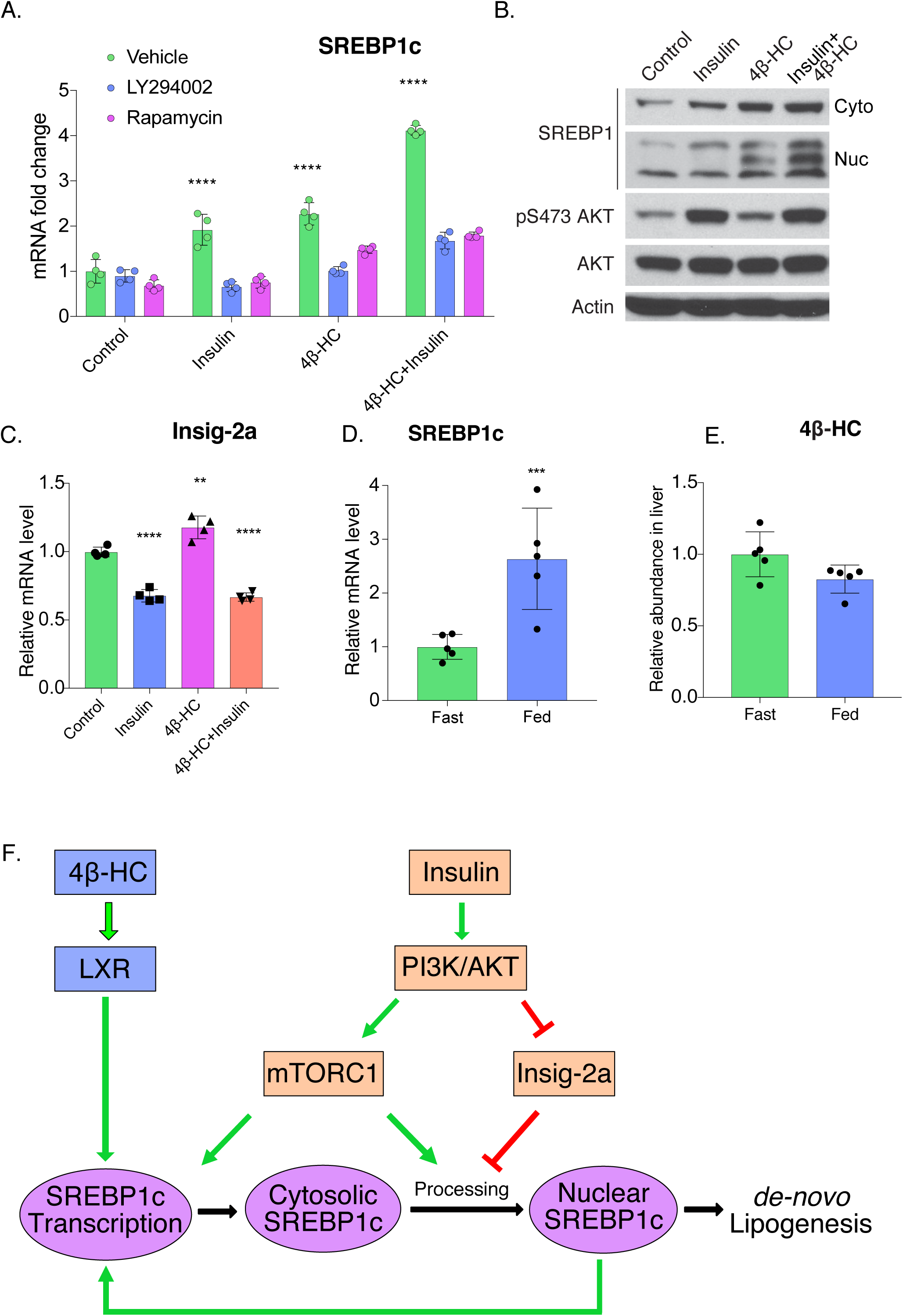
4β-HC acts in parallel to insulin-PI3K signaling to drive SREBP1c expression. **A**. SREBP1c transcription is additive by 4β-HC and Insulin. Primary hepatocytes were treated o/n with vehicle or 5uM 4β-followed with 6hrs stimulation with combinations of Insulin, PI3K inhibitor (LY294002) or Rapamycin. SREBP1c mRNA level was measured by RT-PCR. **B**. 4β-HC and Insulin have an additive effect on SREBP1c expression and nuclear processing. Primary Hepatocytes were treated o/n with vehicle or 4β-HC followed by addition of Insulin for 40min. proteins were extracted and SREBP1 and AKT protein level were measured. **C**. Primary hepatocytes were treated with 4β-HC and Insulin as described in (A) followed by RT-PCR measurement of Insig-2a mRNA level. **D**. Insulin does not induce 4β-HC synthesis. Mice were fasted for 16hrs and then refed for 4 hours, followed by liver extraction and RT-PCR for SREBP1c mRNA level and (**E**.) 4β-HC levels by mass spectrometry. **F**. Model: the 4β-HC-LXR pathway acts in parallel to the insulin-PI3K pathway to drive SREBP1c expression in an additive manner. Bars are Mean+SD. Statistical significance calculated by one-way ANOVA. **p<0.01, ***p<0.001, ****p<0.0001.

To further elucidate the relationship between 4β-HC and insulin, we compared their effects on AKT phosphorylation and SREBP1 protein levels. While insulin strongly increased AKT phosphorylation in primary hepatocytes, we observed a mild increase in SREBP1 expression and processing (Fig 5B). In contrast, 4β-HC stimulation alone was able to induce SREBP1 expression and processing without any change to AKT phosphorylation, suggesting AKT activation is not required for 4β-HC action (Fig 5B). Combined stimulation with insulin and 4β-HC had an additive effect on the levels of both cytoplasmic and nuclear (processed) forms of SREBP1c, supporting our transcriptional data.

Consistent with previous reports, we also detected a marked decrease in Insig-2a mRNA in insulin-stimulated hepatocytes (Fig. 5C). In contrast, 4β-HC caused a mild increase of Insig-2a mRNA levels, and combined insulin and 4β-HC was similar to insulin alone, suggesting Insig-2a downregulation is not required for 4β-HC-dependent SREBP1c activation (Fig. 5C).

To further probe possible connections between insulin-PI3K and 4β-HC-LXR signaling, we tested whether insulin signaling promotes 4β-HC synthesis. Previous reports had shown that in humans 4β-HC has a very slow kinetics, with an extremely long half-life in plasma (60 hours) (Bodin et al., 2002). Pharmacological induction of the main 4β-HC-synthesizing enzyme, cytochrome P450 3A (CYP3A), double 4β-HC concentration in human plasma in 8 days (Kasichayanula et al., 2014), a very different pattern from insulin, which peaks within 1-2 hours following a meals and drops in between. On the other hand, *in vitro* work in primary rat hepatocytes led to the hypothesis that insulin signaling may produce an unknown LXR ligand that, in turn, induces SREBP1c (Chen et al., 2004). To test the possibility of insulin-dependent 4β-HC production, we compared 4β-HC levels in the liver of mice that were either fasted or refed. While mice that were refed showed significant induction of SREBP1c transcription, consistent with SREBP1c regulation by insulin (Fig 5D), the levels of 4β-HC did not increase accordingly (Fig 5E). Collectively, these data suggest that insulin does not induce 4β-HC production according to fasting/feeding cycles, and that 4β-HC most likely acts in parallel to insulin-PI3K signaling in driving SREBP1c transcription and SREBP1c-dependent DNL (Fig 5F).

## Discussion

Here we identify 4β-HC as a unique oxysterol that activates SREBP1c expression and promotes lipogenic gene programs, resulting in induction of fatty acid biosynthesis and cellular accumulation of triglycerides in lipid droplets both in cell culture and *in vivo*. Our results are most consistent with a model in which 4β-HC acts in parallel to insulin-PI3K-mTOR signaling, and the two pathways have additive effects on SREBP1c activation. A simple mechanism that explains the additive effect is that the SREBP1c promoter contains both an LXR binding element (LXRE) and an SREBP binding element (SRE), and transcription can be initiated by the two transcription factors independently and without synergism (Herschlag and Johnson, 1993). Collectively, 4β-HC kinetics suggest that it stimulate SREBP1c expression in a chronic manner, setting the baseline expression, while insulin acts acutely in the postprandial state.

The physiological contexts under which 4β-HC-mediated lipogenesis occurs remain to be determined. Interestingly, several groups using different animal models (mice, rats, rabbits and swine) had all observed that 4β-HC levels increase when animals are fed a high cholesterol diet (Kim et al., 2014; Serviddio et al., 2016; Shimabukuro et al., 2016; Wooten et al., 2014), while a high-fat but with low-cholesterol diet reduces 4β-HC levels in mice (Guillemot-Legris et al., 2016a). Dietary cholesterol was shown to increase SREBP1c expression in an LXR-dependent manner (Peet et al., 1998; Repa et al., 2000). Furthermore, genetically disrupting hepatic cholesterol synthesis through SREBP2 knockout also causes SREBP1c down regulation, which can be rescued by an LXR agonist (Rong et al., 2017). This study also determined that 4β-HC levels are decreases in young SREBP2-null mice, defining a correlation between SREBP2-dependent cholesterol synthesis, 4β-HC levels and SREBP1c expression. Together with this published literature, our results strongly suggest that 4β-HC may be the cholesterol-derived molecule that induces SREBP1c activation via LXR.

An important question is why 4β-HC is the sole oxysterol ligand of LXRs to activate SREBP1 expression in our hands. Several possibilities can be envisioned. The LXR-RXR heterodimer can recruit co-activators (PGC-1*α*, TRRAP, ACS-2, p300, SRC-1) as well as co-repressors (NCoR, SMRT) to the promoters of target genes in a ligand-dependent manner (Hu et al., 2003; Huuskonen et al., 2004; Oberkofler et al., 2003; Wagner et al., 2003; Zhang et al., 2004), but whether all LXR ligands are equally effective in recruiting specific combinations of cofactors is unclear. Supporting this model was an observation in macrophages that the ability of LXR to Recruit RNA polymerase II to SREBP1c promoter requires a specific LXR ligand, while recruitment of RNA polymerase II to the ABCA1 promoter is more promiscuous (Ignatova et al., 2013). Thus, 4β-HC may be able to direct a unique set of co-activators and RNA polymerase II to the SREBP1c promoter, resulting in its activation.

Consistent with previous reports, the synthetic LXR agonist GW3965 was also able to trigger SREBP1 expression (Grefhorst et al., 2002; Schultz et al., 2000). Synthetic LXR agonists are generally more potent than natural LXR ligands, possibly reflecting higher affinity for the ligand-binding site of LXR. By analogy, 4β-HC may bind to LXR with higher affinity than other oxysterol ligands. In turn, higher affinity may translate into longer residence time on the SREBP1c promoter DNA, a possible prerequisite for its efficient activation.

Our data points to the importance of the enzyme that produces 4β-HC, Cyp3A4 (Cyp3A11 in mice) (Bodin et al., 2001), as a crucial regulator of lipogenesis. Consistent with that, several groups have reported that increased Cyp3A4 expression by over expressing its activator, Pregnane X Receptor (PXR), correlated with increases lipogenic gene expression as well as liver triglyceride levels (Huang et al., 2016; Zhou et al., 2006). Conversely, decreased Cyp3A4 expression (He et al., 2013) or its pharmacological inhibition (Chudnovskiy et al., 2014) were associated with lower lipogenic gene expression and liver triglyceride levels. Taken together, these data suggest that Cyp3A4 and 4β-HC may regulate diet-induced lipogenic genes and liver triglyceride levels.

From a more clinical perspective, 4β-HC might have an aggravating effect on the development of Non-Alcoholic Fatty Liver disease (NAFLD). NAFLD is characterized by elevated liver triglycerides not due to alcohol consumption or any other known causes (Neuschwander-Tetri and Caldwell, 2003). Elevated triglyceride levels are associated with LXR and SREBP1c upregulation in NAFLD (Higuchi et al., 2008). NAFLD patients show a significant increase in 4β-HC plasma levels compared to healthy patients (Ikegami et al., 2012). Thus, it is plausible that elevated 4β-HC levels could be an unrecognized driver of triglyceride accumulation in NAFLD. It would be interesting to determine the effect of pharmacologic Cyp3A4 inhibition on disease progression in NAFLD patients

In conclusion, this work highlights a role for 4β-HC, which was long viewed as an ‘orphan’ oxysterol, in regulating lipid metabolism in the liver together with insulin. Future work, dissecting the role of 4β-HC in other organs and in different pathological settings will provide a full picture on the function and significance of this highly abundant oxysterol.

## ACKNOWLEDGEMENTS

We thank all members of the Zoncu Lab for helpful insights. This work was supported by NIH R01GM127763 and R01GM130995 to R.Z., a National Niemann-Pick foundation postdoctoral fellowship to O.M, a National Institutes of Health R01 HL067773 to D.O. and D.C. The Taylor Family Institute for Innovative Psychiatric Research to D.C. Research reported in this publication was supported in part by the National Institutes of Health S10 program under award number 1S10RR026866-01. The content is solely the responsibility of the authors and does not necessarily represent the official views of the National Institutes of Health.

## Material and methods

### Material

Reagents were purchased from the following sources. Antibodies used are: SREBP1 (2A4, Santa cruz), SREBP2 (30682, Abcam), LXRα (PP-PPZ0412-00, R&D systems), LXRβ (K8917, R&D systems), phospho-T308 AKT (C31E5E), AKT (11E7) Cell signaling technologies.

Drugs used are: Cycloheximide (Cell signaling Technology) was used at 10μg/ml. 9-cis-Retinoic acid (Sigma) was used at 50μM. LXR antagonist GSK-2033 (Axon Medchem) was used at 500nM. LXR agonist GW3965 (Fisher Scientific) was used at 500nM. PI3K inhibitor, LY294002 (Cell Signaling Technologies), was used at 10μM. Rapamycin was used at 100nM and received as a gift from David Sabatini. Methyl-beta-cyclodextrin was purchased from Sigma. All Sterols except for custom-synthesized ent-4β-HC (see below) were purchased from Steraloids. C13-glucose was purchased from Cambridge Isotope laboratories.

### Sterol:MbCD pre-complexing

All Sterols were made to 50mM stocks in ethanol. To deliver the sterols to cells, 1.25mM Sterol was complexed with 25mM Methyl-beta-cyclodextrin (MbCD) and vortexed until the solution is clear. Sterols was added media in indicated concentration and incubation time. Control samples were treated by adding the same volume of ethanol to MbCD, which then was delivered to cells in the same corresponding volume.

### Cell culture

Huh7 Cells were maintained on Dulbecco modified Eagle medium (DMEM, 5g/l Glucose. + Glutamine, Gibco) supplemented with 10% FBS (VMR) and p/s (Gibco). Lipid depleted serum (LDS) was made as described in (Goldstein et al., 1983). For assays, on day one; 10^5 cells were plates in 6cm plates. On day two, media was change to 1% LDS, 1g/l glucose DMEM. On Day three; plates were spiked with pre-complexed sterols for indicated times, concentrations and additional compounds.

Primary mouse hepatocytes were purchased from UCSF liver center. Isolation protocol is based on (Li et al., 2010) and adjusted in the following manner. Mice were fasted over night prior to isolation. Hepatocytes were isolated by perfusion protocol (Seglen, 1972) and plated at density of 7×10^5/well on a 6-well collagen coated plates (Corning) in DMEM supplemented with 10% FBS. Once cells adhere, media was replaced to Medium 199 (GIBCO) containing 100nM Dexamethasone (Sigma), 3,3,5-triiodo-L-thyronine (T3, Sigma) and Insulin-Transferrin-Selenium (Gibco). Next day the same media was used without Insulin-transferrin-Selenium to assay Insulin, 4β-HC and inhibitors at indicated times and concentrations.

### Real Time PCR analysis for gene expression

RNA was extracted using RNAesy kit (Qiagen). 1μg of RNA was reverse transcribed using Super Script III (Invitrogen). Quantitative PCR was preformed using SSO advance (BioRad) in Step one Plus (ABI). List of primers is in table 1

### Protein extraction and western blot

Cells were harvested with RIPA buffer supplemented with Phosphatase inhibitor and protease inhibitor (10 mM Tris-Cl (pH 8.0), 1 mM EDTA, 1% Triton X-100, 0.1% sodium deoxycholate, 0.1% SDS, 140 mM NaCl, 10mM Na-PPi, 10mM Na-Beta-glycerophosphate), sonicated with Bioruptor (Diagenode) and normalized using BCA kit (Thermo Scientific).

### Knock down using siRNA

siRNA ON-TARGET plus smart pool against LXRα (cat# L-003413-00-0005), LXRβ (cat# L-003412-02-0005) or non-targeted siRNA ON-TARGETplus Non-targeting Pool (cat# D-001810-10-05) were purchased from Dharamcon. 5μM siRNA was mixed with 5μl Lipofectamine RNAiMAX (life Technologies) in Opti-MEM (Gibco). siRNA is added to Pre-plated Huh7 (10^5 cells/6cm plate) in regular media w/o penicillin streptomycin for 5 hours followed by replacement to regular media for 72 hours.

### Lipid droplet microscopy

Huh7 were plated on a coverslip coated with Fibronectin (Corning) and treated as indicated with sterols and drugs. Cells were fixed with paraformaldehyde and stained with 1ug/ml BODIPY 493/503 for 1 hour. Coverslips were mounted with Vectashield with DAPI (Vector Laboratories) and imaged on a spinning disk confocal system (Andor Revolution on a Nikon Eclipse Ti microscope). BODIPY signal was measured using ImageJ and normalized by the number of nuclei.

### Triglyceride and Cholesterol measurements

Liver samples were powdered with pastel and mortar and lysed in RIPA buffer. Huh7 cells were also harvested in RIPA buffer. 5μl of Samples was used to measure Triglyceride using Triglyceride Infinity (Thermo Fisher) or cholesterol using Amplex red cholesterol measuring kit (Invitrogen) in a clear 96-well sample. BCA kit (Thermo scientific) was used for normalization to protein level. Absorbance and fluorescence was measured by Perkin-Elmer Envision Multi label plate reader.

### C13 incorporation into Triglycerides

Huh7 Cells were seeded at 200K per 6cm plates. Next day, DMEM media with glutamine, containing 5mM C13 glucose (Cambridge Isotope laboratories) and 1% LDS including oxysterols and LXR agonist were added for 24 hours. C12 glucose treated plates were used as reference. Cells were washed twice with ice cold PBS, scraped and pellets were snap-frozen and kept in −80 for later analysis. Lipid extraction and analysis by LC/MS was preformed as described in (Benjamin et al., 2015).

### Husbandry and Diets

All mouse procedures were performed and approved under the University of California, Berkeley Animal Care and Use Committee. 10-week-old C57BL/6J male mice were purchased from the Jackson Laboratory and housed for one week in our facility under standard conditions before experiments were performed. Free access to water and chow (Lab Diets, #3038) was provided throughout this acclimation period. Afterwards, mice were placed on a diet with 50mg/kg/day 4β-HC or LXR agonist GW3965 10mg/kg/day for 7 days. Powdered 10% by kCal fat diet (Research Diets Inc. #D12450J) was used as the base of each treatment food, forming pellets that were dried overnight at room temperature in a laminar flow hood. After seven days, mice were euthanized using CO_2_ and cervical dislocation.

### Cryosectioning and Fluorescent Histochemistry

Liver samples were fixed using 4% (v/v) paraformaldehyde overnight at 4°C. The next day, samples were cryopreserved using sterile-filtered 30% sucrose (w/v) in DPBS (Gibco, 14190-144). After three days, each sample was placed in a 1:1 30% sucrose:Neg-50 (Richard-Allen Scientific) solution and incubated overnight at 4°C. The samples were then frozen on dry ice using undiluted −50°C and stored at −80°C until sectioning. Sequential 20μM thick sections were obtained from each sample using a Leica CM3050S cryostat.

For nuclei and lipid droplet labeling, sectioned tissue was washed three times at room temperature in DPBS for 5 minutes each. Afterward, DPBS containing 10μM BODIPY (Invitrogen, #D3922) was placed on the samples and incubated for 30 minutes at room temperature in the dark. Next, the slides were washed with DPBS twice before incubating in DPBS containing 5μg/mL DAPI (Invitrogen, D1306) for 10 mins at room temperature in the dark. Following DAPI staining, the slides were washed three times in DPBS for 5 minutes each before being mounted using SlowFade Diamond antifade (Invitrogen, #S36972), and sealing with nail polish overnight. Slides were imaged immediately using a Zeiss LSM710 confocal microscope. Images were developed using the IMARIS (Bitplane) image analysis software suite.

### Synthesis of *ent-4β-HC*

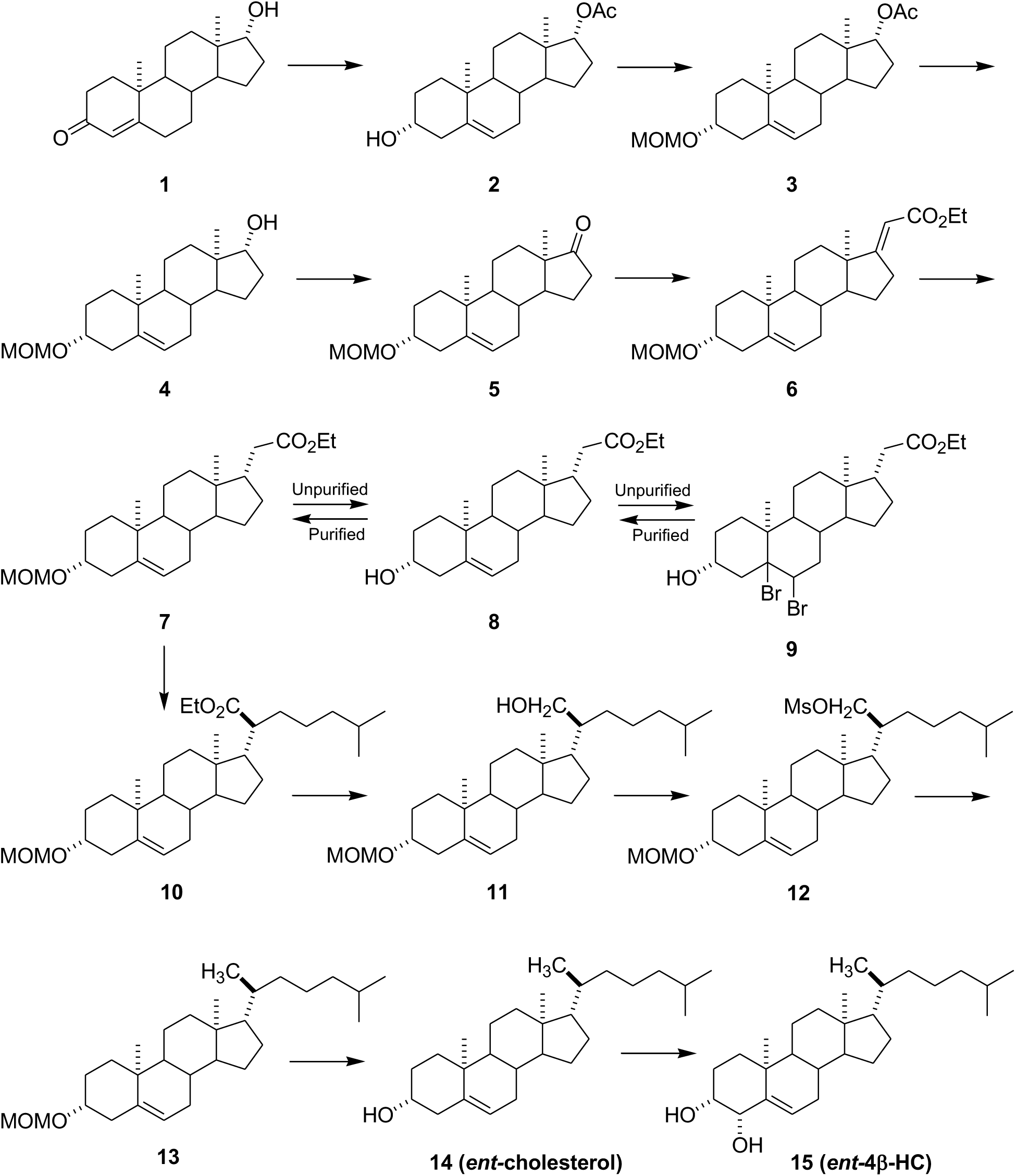

#### *ent-*Steroid 2

*ent-*Testosterone (**1**) was prepared as described previously (Covey, D.F., *Polish J. Chem*., 2006, **80**, 511-522; see also references therein). To a solution of *ent*-testosterone (**1**, 3.8 g, 13.2 mmol) in acetic anhydride (80 mL) was added NaI (7.92 g, 52 mmol) and trimethylsilyl chloride (5.8 mL, 52 mmol) at 0 °C under N_2_. After addition, the reaction was allowed to warm to room temperature for 2 h. The reaction was added to Et_3_N (40 mL) in diethyl ether (100 mL). The ether solution was washed with brine (50 mL x 4), aqueous NaHCO_3_ (50 mL x 2) and dried over Na_2_SO_4_. After filtration, the solvent was removed under reduced pressure and the residue was purified by flash column chromatography (silica gel eluted with 25% EtOAc in hexanes) to give *ent-*steroid **2** (3.05 g, 70%): ^1^H NMR (400 MHz, CDCl_3_) δ 5.33-5.32 (m, 1H), 4.60 (t, *J* = 8.3 Hz, 1H), 3.52-3.47 (m, 1H), 2.30-0.90 (m), 2.02 (s, 3H), 1.00 (s, 3H), 0.79 (s, 3H); ^13^C NMR (100 MHz, CDCl_3_) δ 171.2, 140.9, 121.1, 82.7, 71.5, 51.0, 50.0, 42.3, 42.2, 37.2, 36.7, 36.5, 31.6, 31.5, 31.4, 27.4, 23.5, 21.1, 20.5, 19.3, 11.8.

#### *ent-*Steroid 3

*ent-*Steroid **2** (3.05 g, 4.04 mmol) was dissolved in CH_2_Cl_2_ (50 mL) and cooled to 0 °C. (*i-*Pr)_2_EtN (3.0 mL) and ClCH_2_OMe (1.35 ml, 18.0 mmol) were added and the reaction was stirred at room temperature for 16 h. The reaction was made basic by adding aqueous NaHCO_3_ solution and the product was extracted into CH_2_Cl_2_. The combined extracts were washed with brine, dried over Na_2_SO_4_, filtered and solvent removed to give a viscous liquid which was purified by flash column chromatography (silica gel eluted with 10% EtOAc in hexanes) to give *ent-*steroid **3** as a colorless liquid (2.65 g, 77%): ^1^H NMR (400 MHz, CDCl_3_) δ 5.33-5.32 (m, 1H), 4.65 (s, 2H), 4.59 (t, *J* = 8.2 Hz, 1H), 3.39-3.35 (m, 1H), 3.34 (s, 3H), 2.35-0.89 (m), 2.01 (s, 3H), 0.99 (s, 3H), 0.78 (s, 3H); ^13^C NMR (100 MHz, CDCl_3_) δ 171.0, 140.7, 121.2, 94.6, 82.6, 76.7, 55.0, 50.9, 50.0, 42.3, 39.4, 37.1, 36.7, 31.6, 31.4, 28.8, 27.4, 23.5, 21.0, 20.4, 19.3, 11.8.

#### *ent-*Steroid 4

To a solution of *ent-*steroid **3** (2.65 g, 7.05 mmol) in methanol (60 mL) was added K_2_CO_3_ (4.0 g) at room temperature. The mixture was refluxed for 16 h. Methanol was removed under reduced pressure and the residue was purified by flash column chromatography (silica gel eluted with 25% EtOAc in hexanes) to give *ent-*steroid **4** (2.31 g, 99%): ^1^H NMR (400 MHz, CDCl_3_) δ 5.32-5.30 (m, 1H), 4.64 (s, 2H), 3.61 (t, *J* = 8.6 Hz, 1H), 3.40-3.34 (m, 1H), 3.33 (s, 3H), 2.31-0.87 (m), 0.95 (s, 3H), 0.72 (s, 3H); ^13^C NMR (100 MHz, CDCl_3_) δ 140.7, 121.3, 94.5, 81.6, 76.7, 55.0, 51.2, 50.2, 42.6, 39.4, 37.2, 36.7, 36.5, 31.8, 31.4, 30.3, 28.8, 23.3, 20.5, 19.3, 10.9.

#### *ent-*Steroid 5

To a solution of *ent-*steroid **4** (1.5 g, 4.54 mmol) in CH_2_Cl_2_ (60 mL) was added Dess–Martin periodinane (2.5 g, 6 mmol) at room temperature. After 1 h, water (50 mL) was added, the product was extracted into CH_2_Cl_2_ (150 mL x 3) and the combined extracts were washed with brine (50 mL x 2). The organic layer was dried over Na_2_SO_4_, filtered and the solvents removed. The residue was purified by flash column chromatography (silica gel eluted with 10% EtOAc in hexanes) to give *ent-*steroid **5** (1.5 g, 100%): ^1^H NMR (400 MHz, CDCl_3_) δ 5.39-5.38 (m, 1H), 4.68 (s, 2H), 3.45-3.38 (m, 1H), 3.37 (s, 3H), 2.49-0.98 (m), 1.03 (s, 3H), 0.88 (s, 3H); ^13^C NMR (100 MHz, CDCl_3_) δ 221.0, 140.9, 120.9, 94.7, 76.7, 55.1, 51.7, 50.2, 47.5, 39.5, 37.1, 36.8, 35.8, 31.4, 31.3, 30.8, 28.8, 21.8, 20.3, 19.3, 13.5.

#### *ent-*Steroid 6

A solution of freshly prepared sodium ethoxide (sodium 0.4 g, 15 mmol dissolved in ethanol 15 mL) was added dropwise slowly to a solution of *ent-*steroid **5** (1.5 g, 4.54 mmol) and triethyl phosphonoacetate (3.44 g, 15 mmol) in anhydrous ethanol (25 mL) under N_2_ while stirring at 35-40 °C. After addition, the reaction was refluxed for 16 h. After cooling to room temperature, the ethanol was removed and the residue was dissolved in ether which was washed with water, dried over Na_2_SO_4_ and filtered. Solvent was removed and the residue was purified by flash column chromatography (silica gel eluted with 10% EtOAc in hexanes) to give *ent-*steroid **6** (1.68 g, 87%): ^1^H NMR (400 MHz, CDCl_3_) δ 5.52 (s, 1H), 5.35-5.34 (m, 1H), 4.66 (s, 2H), 4.15-4.09 (m, 2H), 3.43-3.33 (m, 1H), 3.35 (s, 3H), 2.84-2.79 (m, 2H), 2.36-0.93 (m), 1.01 (s, 3H), 0.82 (s, 3H); ^13^C NMR (100 MHz, CDCl_3_) δ 176.1, 167.3, 140.7, 121.3, 108.6, 94.6, 76.7, 59.4, 55.1, 53.7, 50.2, 46.0, 39.5, 37.2, 36.8, 35.1, 31.6, 31.5, 30.4, 28.8, 24.4, 20.9, 19.3, 18.2, 14.3.

The reaction sequence reported below that converts *ent-*steroid **6** into *ent-*steroid **16 (*ent-*VP1-001)** is based on that reported previously for the preparation of the natural stereoisomer of *ent-*steroid **16 (**Wicha, J.; Bal, K. *J. C. S. Perkin I*, **1978**, 1282-1288).

#### Unpurified *ent-*Steroid 7

To a solution of *ent-*steroid **6** (1.4 g, 3.48 mmol) in EtOAc (150 mL) was added PtO_2_ (15 mg) at room temperature. Hydrogenation was carried out under 20 psi for 6 h. Solvent was removed and the residue was purified by flash column chromatography (silica gel eluted with 10% EtOAc in hexanes) to give unpurified *ent-*steroid **7** (1.4 g, 100%): ^1^H NMR δ 4.63-4.60 (m, 1H), 4.08-4.03 (m, 2H), 3.48-3.32 (m, 1H), 3.31 (s, 3H), 2.34-0.57 (m), 0.76 (s, 3H), 0.54 (s, 3H); ^13^C NMR δ 176.1, 140.7, 121.3, 94.4, 76.2, 60.0, 55.3, 55.0, 54.5, 46.9, 44.9, 42.1, 37.4, 37.0, 35.6, 35.5, 35.3, 35.2, 32.1, 28.7, 28.1, 24.4, 20.9, 14.2, 12.5.

Unpurified *ent-*steroid **7** contains minor amounts of the *ent-*steroid in which the Δ^5^ double bond has been hydrogenated. This saturated *ent-*steroid could not be removed easily by chromatography on silica gel. To separate the two compounds chromatographically, *ent-*steroid **7** was converted first into *ent-*steroid **8** and then into *ent-*steroid **9** which is easily purified. *ent-*Steroid **9** was then converted back via *ent-*steroid **8** into *ent-*steroid **7** and then subsequently into *ent-*steroid **10**.

#### Unpurified *ent-*Steroid 8

Acetyl chloride (2 mL) was slowly added to unpurified hydrogenation product *ent-*steroid **7** (1.4 g, 3.48 mmol) in ethanol (30 mL) at room temperature. After 2 h, water was added and the product was extracted into CH_2_Cl_2_ (100 mL x 2). The combined extracts were dried over Na_2_SO_4_, filtered and the solvent was removed under reduced pressure. The residue was purified by flash column chromatography (silica gel eluted with 25% EtOAc in hexanes) to give unpurified *ent-*steroid **8** (1.2 g): ^1^H NMR (400 MHz, CDCl_3_) δ 5.35-5.34 (m, 1H), 4.13-4.07 (m, 2H), 3.55-3.47 (m, 1H), 2.38-0.81 (m), 1.10 (s, 3H), 0.61 (s, 3H); ^13^C NMR (100 MHz, CDCl_3_) δ 173.9, 140.8, 121.5, 71.6, 60.1, 55.5, 50.3, 46.8, 42.2, 41.9, 37.3, 37.2, 36.5, 35.2, 31.9, 31.8, 31.6, 28.1, 24.5, 20.8, 19.4, 14.2, 12.4.

#### *ent-*Steroid 9

To a solution of unpurified *ent-*steroid **8** (1.2 g, 3.33 mmol) in diethyl ether (100 mL) and acetic acid (5 mL) was slowly added Br_2_ in HOAc (3 mL) until a brown color persisted. After 5 min, aqueous Na_2_S_2_O_3_ was added and the reaction became colorless. EtOAc (100 mL) was added and the EtOAc solution was washed with aqueous NaHCO_3_ (50 mL x 2), brine (50 mL) and dried over anhydrous Na_2_SO_4_. After filtration, the solvent was removed under reduced pressure and the residue was purified by flash column chromatography (silica gel eluted with 20% EtOAc in hexanes) to give *ent-*steroid **9** (1.4 g, 81%): ^1^H NMR (400 MHz, CDCl_3_) δ 4.82-4.81 (m, 1H), 4.44-4.37 (m, 1H), 4.12-4.06 (m, 2H), 2.72-1.08 (m), 1.43 (s, 3H), 0.62 (s, 3H); ^13^C NMR (100 MHz, CDCl_3_) δ 173.8, 89.6, 68.9, 60.1, 56.0, 54.0, 47.6, 46.6, 45.6, 42.2, 42.0, 37.2, 37.0, 36.7, 35.2, 30.9, 30.1, 28.0, 24.2, 21.0, 20.3, 14.2, 12.7.

#### Purified *ent-*Steroid 8

Zinc dust (6.0 g) was added to a solution of *ent-*steroid **9** (1.4 g, 2.7 mmol) in HOAc (20 mL) and EtOAc (30 mL) at room temperature. After 16 h, the mixture was filtered through Celite and washed with EtOAc (200 mL). Solvent was removed under reduced pressure and the residue was purified by flash column chromatography (silica gel eluted with 25% EtOAc in hexanes) to give purified *ent-*steroid **8** (925 mg, 95%): ^1^H NMR (400 MHz, CDCl_3_) δ 5.26-5.25 (m, 1H), 4.06-4.01 (m, 2H), 3.85 (s, br, 1H), 3.47-3.40 (m, 1H), 2.31-0.73 (m), 0.93 (s, 3H), 0.54 (s, 3H); ^13^C NMR (100 MHz, CDCl_3_) δ 173.8, 140.7, 121.1, 71.2, 60.0, 55.4, 50.1, 46.6, 41.9, 41.7, 37.1, 37.0, 36.3, 35.0, 31.7, 31.7, 31.2, 27.9, 24.3, 20.6, 19.2, 14.0, 12.2.

#### Purified *ent-*Steroid 7

Purified *ent-*steroid **8** (925 mg, 2.57 mmol) was dissolved in CH_2_Cl_2_ (20 mL) and cooled to 0 °C. (*i-*Pr)_2_EtN (1.3 mL, 7.5 mmol) and ClCH_2_OMe (0.45 ml, 6.0 mmol) were added and the reaction was stirred at room temperature for 16 h. The reaction mixture was made basic by adding aqueous saturated NaHCO_3_ solution and the product extracted into CH_2_Cl_2_. The combined extracts were washed with brine, dried over anhydrous Na_2_SO_4_ and solvent removed to give a viscous liquid which was purified by flash column chromatography (silica gel eluted with 20% EtOAc in hexanes) to give purified *ent-*steroid **7** as a colorless liquid (1.02 g, 98%): ^1^H NMR (400 MHz, CDCl_3_) δ 5.34-5.33 (m, 1H), 4.67 (s, 2H), 4.12 (q, *J* = 7.0 Hz, 2H), 3.42-3.36 (m, 1H), 3.35 (s, 3H), 2.37-0.80 (m), 1.00 (s, 3H), 0.60 (s, 3H); ^13^C NMR (CDCl_3_) δ 173.8, 140.7, 121.5, 94.6, 76.8, 60.0, 55.5, 55.1, 50.3, 46.7, 41.9, 39.5, 37.2, 37.1, 36.7, 35.2, 31.9, 31.8, 28.9, 28.1, 24.5, 20.7, 19.3, 14.2, 12.3.

#### *ent-*Steroid 10

To a solution of the *ent-*steroid **7** (202 mg, 0.5 mmol) in THF (10 mL) was added LDA (0.75 mL, 2.0 M in THF, 1.5 mmol) and HMPA (0.29 mL, 1.65 mmol) at –78 °C. After 1 h, 1-bromo-4-methylpentane (0.44 mL, 3 mmol) was added. After addition, the reaction was warmed to room temperature for 16 h. Aqueous NH_4_Cl was added and extracted with EtOAc (100 mL x 2) and the combined extracts were dried over anhydrous Na_2_SO_4_. Solvent was removed under reduced pressure and the residue was purified by flash column chromatography (silica gel eluted with 20% EtOAc in hexanes) to give *ent-*steroid **10** (236 mg, 97%): ^1^H NMR (400 MHz, CDCl_3_) δ 5.34-5.33 (m, 1H), 4.67 (s, 2H), 4.13-4.08 (q, *J* = 7.4 Hz, 2H), 3.41-3.37 (m, 1H), 3.35 (s, 3H), 2.35-0.79 (m), 0.98 (s, 3H), 0.70 (s, 3H); ^13^C NMR (100 MHz, CDCl_3_) δ 176.2, 140.7, 121.5, 94.6, 76.9, 59.6, 56.0, 55.1, 52.6, 50.1, 47.4, 41.9, 39.5, 38.8, 37.5, 37.2, 36.7, 32.2, 31.8, 31.7, 28.9, 27.8, 27.0, 25.0, 23.8, 22.7, 22.3, 20.8, 19.3, 14.2, 12.0.

#### *ent-*Steroid 11

To a solution of *ent-*steroid **10** (236 mg, 0.5 mmol) in diethyl ether (20 mL) was added LiAlH_4_ (2.0 M in diethyl ether, 4.0 mL, 8.0 mmol) at room temperature. After 2 h, water (0.32 mL), 10 % of NaOH (0.64 mL) and water (0.96 mL) were slowly added sequentially. After stirring for 30 min, the mixture was filtered through Celite and washed with CH_2_Cl_2_ (100 mL). Solvent was removed under reduced pressure and the residue was purified by flash column chromatography (silica gel eluted with 25% EtOAc in hexanes) to give *ent-*steroid **11** (212 mg, 98%): ^1^H NMR (400 MHz, CDCl_3_) δ 5.34-5.33 (m, 1H), 4.66 (s, 2H), 3.71-3.61 (m, 2H), 3.44-3.36 (m, 1H), 3.34 (s, 3H), 2.35-0.88 (m), 0.99 (s, 3H), 0.68 (s, 3H); ^13^C NMR (100 MHz, CDCl_3_) δ 140.6, 121.6, 94.6, 76.7, 62.5, 56.6, 55.1, 50.3, 50.1, 42.3, 42.0, 39.5, 39.1, 37.2, 36.6, 31.8, 29.5, 28.8, 27.9, 27.5, 24.1, 24.0, 22.7, 22.5, 21.0, 19.3, 12.1.

#### *ent-*Steroid 12

To a solution of *ent-*steroid **11** (212 mg, 0.48 mmol) in CH_2_Cl_2_ (10 mL) was added mesyl chloride (1 mmol, 0.08 mL) and Et_3_N (0.28 mL, 2 mmol) at 0 °C. After 1 h, aqueous NH_4_Cl was added and the product was extracted into CH_2_Cl_2_ (100 mL x 2). The combined extracts were dried over anhydrous Na_2_SO_4_, filtered and the solvents removed. The residue was purified by flash column chromatography (silica gel eluted with 10% EtOAc in hexanes) to give *ent-*steroid **12** (241 mg, 97%): ^1^H NMR (400 MHz, CDCl_3_) δ 5.33-5.32 (m, 1H), 4.66 (s, 2H), 4.36-4.32 (m, 1H), 4.18-4.09 (m, 1H), 3.42-3.37 (m, 1H), 3.34 (s, 3H), 2.97 (s, 3H), 2.34-0.89 (m), 0.98 (s, 3H), 0.69 (s, 3H); ^13^C NMR (100 MHz, CDCl_3_) δ 140.6, 121.4, 94.6, 76.8, 70.0, 56.4, 55.1, 50.0, 49.9, 42.0, 39.7, 39.4, 39.2, 39.0, 37.2, 37.1, 36.6, 31.7, 31.6, 29.4, 28.8, 27.7, 27.4, 24.0, 23.4, 22.6, 22.4, 20.9, 19.3, 12.1.

#### *ent-*Steroid 13

To a solution of *ent-*steroid **12** (241 mg, 0.46 mmol) in diethyl ether (30 mL) was added LiAlH_4_ (2.0 M in diethyl ether, 4.0 mL, 8.0 mmol) at room temperature. After 2 h, water (0.32 mL), 10 % of NaOH (0.64 mL) and water (0.96 mL) were slowly added sequentially. After stirring for 30 min, the mixture was filtered through Celite and washed with CH_2_Cl_2_ (100 mL). Solvent was removed under reduced pressure and the residue was purified by flash column chromatography (silica gel eluted with 10% EtOAc in hexanes) to give *ent-*steroid **13** (188 mg, 95%): ^1^H NMR (400 MHz, CDCl_3_) δ 5.35-5.34 (m, 1H), 4.68 (s, 2H), 3.46-3.38 (m, 1H), 3.36 (s, 3H), 2.37-0.86 (m), 1.01 (s, 3H), 0.68 (s, 3H); ^13^C NMR (100 MHz, CDCl_3_) δ 140.7, 121.7, 94.6, 76.9, 56.7, 56.1, 55.1, 50.1, 42.3, 39.8, 39.5, 39.4, 37.2, 36.7, 36.2, 35.8, 31.9, 31.8, 28.9, 28.2, 28.0, 24.3, 23.8, 22.8, 22.5, 21.0, 19.3, 18.7, 11.8.

#### *ent-*Steroid 14 (*ent-*cholesterol)

To a solution of *ent-*steroid **13** (188 mg, 0.44 mmol) in THF (20 mL) was added 6 N HCl (10 mL) at room temperature. After 4 h, the product was extracted into CH_2_Cl_2_ (100 mL x 2) and the combined extracts were washed with aqueous NaHCO_3_ (50 ml x 2), dried over anhydrous Na_2_SO_4_, and filtered. Solvent was removed under reduced pressure and the residue was purified by flash column chromatography (silica gel eluted with 20% EtOAc in hexanes) to give *ent-*steroid **14** (165 mg, 98%); ^1^H NMR (400 MHz, CDCl_3_) δ 5.36-5.35 (m, 1H), 3.57-3.49 (m, 1H), 2.33-0.86 (m), 1.01 (s, 3H), 0.68 (s, 3H); ^13^C NMR (100 MHz, CDCl_3_) δ 140.7, 121.7, 71.8, 56.7, 56.1, 50.1, 42.3, 42.2, 39.8, 39.5, 37.2, 36.5, 36.2, 35.8, 31.9(2C), 31.6, 28.2, 28.0, 24.3, 23.8, 22.8, 22.6, 21.1, 19.4, 18.7, 11.8.

#### *ent-*Steroid 15 (*ent-*4β-HC)

A procedure previously reported to convert cholesterol to 4β-hydroxycholesterol was used (Nury, T; Samadi, M; Zarrouk, A; Riedinger, J. M; Lizard, G. *Eur. J. Med. Chem*. **2013**, *70*, 558-567.) to convert *ent-* cholesterol **14** into *ent-*4β-hydroxycholesterol **15**.

To a solution of *ent*-cholesterol **14** (29 mg, 0.0747 mmol) in dioxane (5 mL) and water (2 drops) was added SeO_2_ (17 mg, 0.15 mmol) at room temperature. The mixture was heated to 90 °C for 16 h. After cooling to room temperature, solvent was removed under reduced pressure. The residue was purified by flash column chromatography (silica gel eluted with 30% EtOAc in hexanes) to give *ent*-4β-hydroxycholesterol **15** (17 mg, 58%): mp 169-171 °C; [*α*]_D_^20^ +41.7 (*c* = 0.12, CHCl_3_); ^1^H NMR (400 MHz, CDCl_3_) δ 5.69-5.68 (m, 1H), 4.15-4.14 (m, 1H), 3.58-3.55 (m, 1H), 2.20-0.78 (m), 1.19 (s, 3H), 0.69 (s, 3H); ^13^C NMR (100 MHz, CDCl_3_) δ 142.7, 128.8, 77.3, 72.5, 56.9, 56.1, 50.2, 42.3, 39.7, 39.5, 36.9, 36.2, 36.0, 35.8, 32.1, 31.8, 28.2, 28.0, 25.4, 24.2, 23.8, 22.8, 22.5, 21.0, 20.5, 18.7, 11.8; IR (film, cm^-1^) 3406, 1455, 1366, 1072.

